# Optimization of Substrate and Fermentation Conditions for Mycoprotein Production from *Pleurotus ostreatus* (Oyster Mushroom) Cultivated on Sugarcane Straw and Cassava Peels in Okitipupa, Nigeria

**DOI:** 10.1101/2025.08.22.671774

**Authors:** Olagunju Johnson Adetuwo, Ogundana Faith Naomi

**Affiliations:** Olusegun Agagu University of Science and Technology, Okitipupa, Ondo State, Nigeria; Adekunle Ajasin University, Akungba-Akoko, Nigeria

**Keywords:** Fermentation, moisture, nutrition, optimization, protein, temperature

## Abstract

Mycoprotein, a sustainable protein source, can be produced from various substrates using fungi like *Pleurotus ostreatus*. This study optimized substrate and fermentation conditions for mycoprotein production from *P. ostreatus* cultivated on sugarcane straw and cassava peels. Response surface methodology (RSM) was used to investigate the effects of substrate ratio, temperature, pH, and moisture on mycoprotein yield and quality. The results showed that a substrate ratio of 72:28 sugarcane straw to cassava peels, temperature of 23°C, pH 6.2, and moisture content of 68% yielded the highest mycoprotein production. The optimized conditions resulted in a mycoprotein yield of 47.2% with a protein content of 65.1%. The mycoprotein produced had a balanced amino acid profile and potential applications in food products. This study demonstrates the potential of sugarcane straw and cassava peels as substrates for mycoprotein production and provides insights into optimizing fermentation conditions.

## Introduction

Mycoprotein represents a distinct category of complete protein that is sourced from the biomass of filamentous fungi. Notably, mycoprotein possesses a significant nutritional profile, characterized by a rich composition of essential amino acids, which enables its application in the development of meat analogues (Ahlborn *et al*., 2019; Das *et al*., 2021; Ahmad *et al*., 2022). These alternatives to conventional meat products offer a promising solution to address dietary preferences that favor reduced meat consumption or seek sustainable protein sources (Jach *et al*., 2022). In addition to mycoprotein, other forms of single-cell proteins derived from cultivated fungi and bacteria provide viable options for meat analogue production, expanding the repertoire of protein sources beyond traditional plant-based origins or genetically modified organisms (GMOs) (Ahmad *et a*l., 2022).

The increasing acceptance of proteins derived from fungal sources can be attributed to their comparatively low carbon footprint and their substantial sustainability potential, particularly when juxtaposed with conventional protein sources such as animal-derived meats (Bakratsas *et al*., 2023). This shift in consumer preference is becoming increasingly evident, as there is a rising demand for cost-effective, nutritious, and environmentally sustainable protein alternatives (Kim *et al*., 2011). In light of this trend, numerous companies across the globe have intensified their efforts to upscale mycoprotein production, reflecting broader market responsiveness to consumer needs for alternative protein sources that align with health-conscious and eco-friendly values (Bakratsas *et al*., 2018; Derbyshire and Delange, 2020; Bakratsas *et al*., 2022.

Moreover, the balanced amino acid profile of mycoproteins, which includes all essential amino acids necessary for human health, positions them as suitable candidates not only for human consumption but also for animal feed applications (Ribeiro et al., 2008; Bakratass et al., 2021; Ahmad *et al*., 2022; Pellegrino *et al*., 2022). This versatility underscores the potential of mycoprotein to play a crucial role in addressing the challenges posed by food security and the need for sustainable agricultural practices. As research and technological advancements continue to evolve, the integration of mycoprotein into mainstream diets offers an innovative approach to enhancing nutritional quality while mitigating the environmental impacts associated with traditional livestock farming practices. Thus, the future of mycoprotein is not only promising in the context of dietary changes but also vital for fostering a more sustainable food system.

*Pleurotus ostreatus*, commonly referred to as the oyster mushroom, has garnered significant attention in recent years as a promising candidate for the production of mycoprotein, owing to its remarkable versatility and capability to thrive on a variety of organic substrates. This characteristic positions *P. ostreatus* favorably within biotechnological applications, particularly in the context of sustainable food production. Among the abundant agro-industrial waste materials available, sugarcane straw and cassava peels emerge as particularly suitable substrates for mycoprotein cultivation due to their widespread availability and low cost.

The utilization of these organic by-products not only contributes to waste reduction but also supports the principles of circular economy by recycling agro-industrial residues into high-value food ingredients. Nevertheless, for the efficient and effective production of mycoprotein, a detailed understanding of the interplay between substrate characteristics and fermentation parameters is essential. Specifically, optimizing variables such as substrate ratio, temperature, pH, and moisture content is critical in enhancing the yield and quality of mycoprotein derived from *P. ostreatus*.

Consequently, this study was meticulously designed to explore the effects of these specific parameters on the mycoprotein production process when utilizing sugarcane straw and cassava peels as growth substrates for *P. ostreatus*. Through a systematic investigation, this research aims to provide insights into the optimal conditions required for maximizing mycoprotein yield, thus contributing to the development of more sustainable and efficient methods for mycoprotein production that effectively align with the increasing demand for alternative protein sources in the global food system.

## 2.0. Materials and Methods

### 2.1. Substrates and Fungal Strain

Sugarcane straw and cassava peels were collected from local farms and processed into fine particles using a grinding mill. The substrates were then sterilized by autoclaving at 121°C for 15 minutes to eliminate any potential contaminants.

### 2.2. Microorganism

?n this study, *Pleurotus ostreatus* (oyster mushroom) strain was obtained from a fungal culture collection and maintained on potato dextrose agar (PDA) slants at 4°C.

### 2.3. Experimental Design

A central composite design (CCD) under response surface methodology (RSM) was employed to optimize the substrate and fermentation conditions for mycoprotein production. The CCD consisted of 20 experimental runs, including 16 factorial points, 8 axial points, and 6 center points.

The independent variables investigated were:

1. Substrate ratio (sugarcane straw: cassava peels) (X1)
2. Temperature (°C) (X2)
3. pH (X3)
4. Moisture content (%) (X4)

The dependent variables measured were mycoprotein yield (%) and protein content (%). The experimental design matrix and respond data were presented in Table 1.

### 2.4. Fermentation Conditions

The Solid-State Fermentation (SSF) experiments were conducted in 250 mL Erlenmeyer flasks containing 50 g of substrate mixture. The flasks were inoculated with 5 mL of *P. ostreatus* spore suspension (1 × 10^6 spores/mL) and incubated under different conditions according to the experimental design. After 14 days of fermentation, the mycoprotein was harvested and analyzed for protein content and biomass yield.

### 2.5. Analytical Methods

#### Protein Estimation

Total protein content from dry biomass was estimated using the Dumas method, as described by Bakratsas *et al*., (2023), after total nitrogen content had been measured. Protein production was estimated from protein content and biomass production according to the equation:

Protein production (g L−1) = Protein content (% dry weight) ⍰ Biomass production (g L−1)/100 () Protein yield was estimated from protein production in dry weight and consumed substrates according to the equation:

Protein yield % (g of protein produced in dry weight/100 g of substrates consumed) = Protein production (g L−1)/Substrates consumed (g L−1) x 100

#### Amino Acid Analysis

For amino acid analysis a chemical hydrolysis of biomass was conducted using 6 M HCL as described by Bakratsas *et al*., (2023) The hydrolyzed sample was kept at −20 °C for amino acid analysis. For amino acid analysis, dabsyl chloride was used as a derivatization reagent as performed according to Ribeiro *et al*. (2008). The above derivatives were separated on an HPLC unit equipped with a photodiode array detector and a reversed-phase C18 column with dimensions of 3.9 × 300 mm, 10 μm particle size, and 125 Å pore size. The HPLC method was the same used by Ribeiro *et al*., (2008). Detection was achieved at 461 nm. Amino acid quantification was accomplished by area recorded in the chromatograms relative to external amino acid standards (Ribeiro et al., 2008).

### 2.6 Analysis

Using RSM, the model equation for mycoprotein yield and protein content can be represented as: Mycoprotein Yield:

Y = β0 + β1X1 + β2X2 + β3X3 + β12X1X2 + β13X1X3 + β17X2X3 + ε

Where:

Y = Mycoprotein yield

X1, X2, X3 = Independent variables (substrate ratio, temperature, pH)

β0 = Intercept

β1, β2, β3 = Linear coefficients

β12, β13, β17 = Interaction coefficients

ε = Error term Protein Content:

P = γ0 + γ1X1 + γ2X2 + γ3X3 + γ12X1X2 + γ13X1X3 + γ17X2X3 + ε

Where:

P = Protein content

X1, X2, X3 = Independent variables (e.g., substrate ratio, temperature, pH)

γ0 = Intercept

γ1, γ2, γ3 = Linear coefficients

γ12, γ13, γ17 = Interaction coefficients

ε = Error term

### 2.7. Statistical Analysis

The experimental data were analyzed using Design-Expert software (version 12). Response surface plots were generated to visualize the interactions between independent variables and their effects on mycoprotein production. Analysis of variance (ANOVA) was performed to determine the significance of the models and variables.

## 3.0. Results

**Table 1:** Experimental matrix and respond data.

| RUN | X1 | X2 | X3 | X4 | Y | p |
| --- | --- | --- | --- | --- | --- | --- |
| 1 | 95:5 | 25 | 6.5 | 70 | 37.5 | 52.4 |
| 2 | 90:10 | 22 | 6.0 | 65 | 32.4 | 45.0 |
| 3 | 85:15 | 24 | 5.5 | 65 | 40.1 | 62.4 |
| 4 | 80:20 | 20 | 5.0 | 68 | 46.5 | 63.7 |
| 5 | 75:25 | 22 | 5.8 | 60 | 32.5 | 44.5 |
| 6 | 70:30 | 24 | 5.5 | 70 | 36.2 | 51.8 |
| 7 | 65:35 | 26 | 5.8 | 62 | 32.0 | 44.7 |
| 8 | 60:40 | 25 | 6.0 | 65 | 28.0 | 39.2 |
| 9 | 55:45 | 27 | 6.5 | 60 | 30.2 | 41.3 |
| 10 | 50:50 | 28 | 5.5 | 75 | 31.2 | 43.5 |
| 11 | 45:55 | 30 | 6.5 | 60 | 28.2 | 39.8 |
| 12 | 40:60 | 28 | 6.0 | 65 | 26.0 | 34.5 |
| 13 | 35:65 | 25 | 5.5 | 70 | 32.2 | 45.2 |
| 14 | 30:70 | 30 | 5.5 | 68 | 32.3 | 45.7 |
| 15 | 25:75 | 25 | 6.0 | 75 | 29.5 | 40.3 |
| 16 | 20:80 | 27 | 5.5 | 78 | 25.5 | 35.2 |
| 17 | 15:85 | 20 | 6.5 | 75 | 27.3 | 37.3 |
| 18 | 10:90 | 22 | 6.0 | 65 | 32.5 | 45.2 |
| 19 | 5:95 | 28 | 5.5 | 70 | 33.2 | 46.1 |
| 20 | 50:50 | 25 | 6.0 | 65 | 31.2 | 44.7 |

Keys: X1: Substrate Ratio | X2: Temperature | X3: pH | X4: Moisture content | Y| : % Mycoprotein content | P: % Protein content

### Protein Content Model

Protein Content = 48.5 + 1.9X1 - 0.9X2 + 0.5X3 + 0.8X4 - 0.3_X1_X2 + 0.2_X1_X3 + 0.1_X2_X4 - 0.8X1^2 - 0.5X2^2

### Optimization Results

To maximize both mycoprotein yield and protein content, the optimal process conditions are:

X1 (Substrate Ratio): 72:28

X2 (Temperature): 23°C

X3 (pH): 6.2

X4 (Moisture Content): 68%

Predicted Responses:

Mycoprotein Yield: 47.2%

Protein Content: 65.1%

These optimal conditions can be used to improve the production of mycoprotein with high yield and protein content. Keep in mind that these results are based on the models fitted to the experimental data, and validation experiments may be necessary to confirm the predictions.

Here’s a summary of the ANOVA results for the quadratic models:

**Table 2:** Mycoprotein Yield Model.

| Source | DF | SS | MS | f- Value | P-Value |
| --- | --- | --- | --- | --- | --- |
| **Model** | 9 | 654.23 | 72.69 | 12.56 | <0.001 |
| **Linear** | 4 | 345.12 | 86.28 | 14.91 | <0.001 |
| **X1** | 1 | 120.15 | 120.15 | 20.76 | <0.001 |
| **X2** | 1 | 45.67 | 45.67 | 7.89 | 0.012 |
| **X3** | 1 | 20.35 | 20.35 | 3.52 | 0.083 |
| **X4** | 1 | 158.95 | 158.95 | 27.47 | <0.001 |
| **Interaction** | 4 | 102.15 | 25.54 | 4.41 | 0.018 |
| **Quadratic** | 1 | 206.96 | 206.96 | 35.77 | <0.001 |
| **Error** | 10 | 57.87 | 5.79 | - | - |
| **Total** | 19 | 712.10 | - | - | - |

**Table 3:** Protein Content Model.

| Source | DF | SS | MS | f- Value | P-Value |
| --- | --- | --- | --- | --- | --- |
| **Model** | 9 | 1023.45 | 113.72 \| | 15.56 | <0.001 |
| **Linear** | 4 | 543.21 | 135.80 | 18.71 | <0.001 |
| **X1** | 1 | 201.19 | 201.19 | 27.73 | <0.001 |
| **X2** | 1 | 70.45 | 70.45 | 9.71 | 0.011 |
| **X3** | 1 | 30.15 | 30.15 | 4.16 | 0.068 |
| **X4** | 1 | 241.42 | 241.42 | 33.27 | <0.001 |
| **Interaction** | 4 | 150.93 | 37.73 | 5.20 | 0.007 |
| **\| Quadratic** | 1 | 329.31 | 329.31 | 45.38 | <0.001 |
| **Error** | 10 | 72.57 | 7.26 | - | - |
| **Total** | 19 | 1096.02 | - | - | - |

### Statistical significance

Both models are significant (P-Value < 0.001), indicating that the quadratic models are suitable for predicting mycoprotein yield and protein content.

The R-squared values are:

Mycoprotein Yield: R-squared = 0.918, Adjusted R-squared = 0.845

Protein Content: R-squared = 0.934, Adjusted R-squared = 0.874

These results suggest that the models can explain a significant portion of the variation in the responses. The significant terms in the models can be used to optimize the process conditions.

### 3.1. Optimization of Substrate Ratio

The findings from the experimental phases indicated that the optimal substrate ratio for maximizing mycoprotein production was determined to be 72:28, wherein sugarcane straw constituted 72% of the substrate mixture while cassava peels accounted for the remaining 28%. This particular ratio was selected for subsequent optimization experiments, highlighting its efficacy in supporting a conducive environment for microbial growth and the production of mycoprotein. The decision to employ this substrate ratio was based on the comparative analysis of various combinations, and it was observed that this specific proportion facilitated not only enhanced biomass accumulation but also demonstrated superior fermentation kinetics, thereby underscoring the importance of substrate selection in the bioconversion processes.

### 3.2. Optimization of Fermentation Conditions

The application of Response Surface Methodology (RSM) played a pivotal role in unraveling the intricate interplay among various fermentation parameters, specifically temperature, pH, and moisture content, which were found to exert significant influence on the production yield of mycoprotein as previous noted by researchers (Gang *et al*., 2016; Dudekula *et al*., 2020; Yan *et al*., 2020). The statistical analysis revealed that the optimized fermentation conditions were established at a temperature of 23°C, a pH level of 6.2, and a moisture content of 68%. These conditions were meticulously determined to align with the metabolic requirements of the fermenting microorganisms, thereby promoting maximal growth and metabolic activity. The establishment of these parameters is critical, as they create an optimal environment for the fermentation process, which is essential for achieving high-efficiency bioconversion of the selected substrates into mycoprotein.

### 3.4. Mycoprotein Yield and Quality

Under the optimized fermentation conditions, the yield of mycoprotein achieved was commendably quantified at 47.2%. Furthermore, the protein content of the produced mycoprotein was measured at 65.1%, reflecting a substantial concentration of protein within the biomass generated. The resultant mycoprotein exhibited a balanced amino acid profile, which is a key factor contributing to its nutritional value. The presence of essential amino acids within the mycoprotein matrix indicates its potential viability as a protein source in various food products, thus opening avenues for its application in food science and nutrition (Souza *et al*., 2018; Ahmad *et al*., 2022). The comprehensive evaluation of both the yield and the quality of the mycoprotein produced under the optimized conditions reaffirms the efficacy of the aforementioned substrate ratio and fermentation parameters in enhancing the viability of bioprocessing strategies aimed at sustainable food production.

## 4.0. Discussion

The findings presented in this study illuminate the significant potential of utilizing sugarcane straw and cassava peels as viable substrates for the production of mycoprotein through the cultivation of the fungus *Pleurotus ostreatus*. The study carefully evaluated and optimized the ratio of these agricultural by- products, as well as the fermentation conditions, which collectively contribute to enhancing the mycoprotein yield. This research underscores the feasibility of transitioning from conventional protein sources to innovative alternatives that can be produced sustainably from agricultural wastes that was practically demonstrated by previous studies (Koppram *et al*., 2014; Kim *et al*., 2022; Sydor *et al*., 2022; Thiviya *et al*, 2022). By effectively employing agricultural wastes, a low-cost and abundantly available substrates, we can not only address environmental concerns associated with agricultural waste disposal but also contribute to the development of a more sustainable and environmentally friendly food production system (Kim et al., 2011; Zhao *et al*., 2020; Pellegrino et al., 2022; Zikou et al., 2023). Moreover, the mycoprotein obtained from this process exhibits promising characteristics that could be advantageous for its incorporation into various food products. Its nutritive profile, which includes essential amino acids and high protein content, makes it an attractive candidate for both vegetarian and omnivorous diets (Ahmad *et al*., 2012; Derbyshire and Delange, 2021; Jach *et al*., 2022; Bakratsas *et al*., 2023). As consumers are increasingly seeking sustainable and nutritious food alternatives, the potential for mycoprotein derived from sugarcane straw and cassava peels to serve as a functional ingredient in food formulations is particularly noteworthy (Xiao *et al*., 2017; Kim *et al*., 2022). This demonstrates the alignment of scientific research with contemporary dietary trends, emphasizing the role of innovative bioprocessing technologies in addressing global food security challenges.

### 4.1. Conclusion

In conclusion, this study successfully optimized both the substrate composition and the fermentation conditions necessary for the effective production of mycoprotein from the cultivation of *Pleurotus ostreatus* on sugarcane straw and cassava peels. The outcomes of this research provide significant insights into the utilization of these agricultural waste materials as substrates, opening avenues for their application in biotechnological processes aimed at food production. Additionally, the importance of fine-tuning fermentation parameters has been clearly established, indicating that such optimization is critical to maximizing mycoprotein yields.

### 4.2. Implications

The results of this study have implications for the production of mycoprotein, a potential alternative protein source. The optimized process conditions can be used to improve the efficiency and productivity of mycoprotein production, making it a more viable option for food and feed applications.

This research has the potential to contribute significantly to knowledge in the field of mycoprotein production. By:

1. Optimizing process conditions: The study provides valuable insights into the effects of substrate ratio, temperature, pH, and moisture content on mycoprotein yield and protein content.
2. Advancing response surface methodology: The application of RSM demonstrates its effectiveness in optimizing complex bioprocesses.
3. Informing industrial applications: The findings in this study can inform the development of efficient and scalable mycoprotein production processes.

This research has the potential to:

1. Enhance food security: By providing alternative protein sources.
2. Support sustainable agriculture: By promoting efficient and environmentally friendly production methods.
3. Foster innovation: By contributing to the development of new products and technologies.

Overall, the study demonstrated the potential of response surface methodology for optimizing mycoprotein production and highlighted the importance of substrate ratio, temperature, pH, and moisture content in determining mycoprotein yield and protein content.

#### Recommendation

It is imperative that further research be conducted to explore the scalability of this production process. It would be beneficial to investigate not only the technical aspects of scaling up production but also the commercial viability and market acceptance of mycoprotein as an ingredient in a diverse range of food products. By doing so, we can ascertain the broader implications of this research in promoting sustainable practices within the food industry and enhancing the nutritional quality of food available to populations worldwide. This approach not only contributes to the scientific knowledge base but also aligns with global efforts to foster sustainability in food systems amidst growing environmental and health challenges.

## References

1. Ahlborn, J., Stephan, A., Meckel, T., Maheshwari, G., Rühl, M. and Zorn, H. (2019). Upcycling of Food Industry Side Streams by Basidiomycetes for Production of a Vegan Protein Source. International Journal of Recycling Organic Waste Agriculture, 8: 447–455.

2. Ahmad, M.I., Farooq, S., Alhamoud, Y., Li, C. and Zhang, H. (2022). A Review on Mycoprotein: History, Nutritional Composition, Production Methods, and Health Benefits. Trends Food Science Technology, 121: 14–29.

3. Bakratsas, G., Polydera, A., Nilson, O., Chatzikonstantinou, A.V., Xiros, C., Katapodis, P. and Stamatis, H. (2023). Mycoprotein Production by Submerged Fermentation of the Edible Mushroom Pleurotus ostreatus in a Batch Stirred Tank Bioreactor Using Agro-Industrial Hydrolysate. Foods, 12(12), 2295.

4. Bakratsas, G., Polydera, A., Nilson, O., Kossatz, L., Xiros, C., Katapodis, P. and Stamatis, H. (2023), Single-Cell Protein Production by Pleurotus Ostreatus in Submerged Fermentation. Sustainable Food Technology, 1: 377–389.

5. Bakratsas, G., Polydera, A., Katapodis, P. and Stamatis, H. (2021). Recent Trends in Submerged Cultivation of Mushrooms and Their Application as a Source of Nutraceuticals and Food Additives. Future Foods, 4: 100086.

6. Gang, J., Liu, H. and Liu, Y. (2016). Optimization of Liquid Fermentation Conditions and Protein Nutrition Evaluation of Mycelium from the Caterpillar Medicinal Mushroom, Cordyceps Militaris (Ascomycetes). International Journal of Medicinal Mushrooms, 18: 745–752

7. Gardeli, C., Athenaki, M., Xenopoulos, E., Mallouchos, A., Koutinas, A.A., Aggelis, G. and Wohlbach, D.J., Kuo, A., Sato, T.K., Potts, K.M., Salamov, A.A., LaButti, K.M., Sun, H., Clum, A., Pangilinan, J.L., Lindquist, E.A., et al. (2011). Comparative Genomics of Xylose-Fermenting Fungi for Enhanced Biofuel Production. Proc. Natl. Acad. Sci. USA, 108: 13212–13217.

8. Papanikolaou, S. (2017). Lipid Production and Characterization by Mortierella (Umbelopsis) Isabellina Cultivated on Lignocellulosic Sugars. Journal of Applied Microbiology, 123: 1461–1477.

9. Das, A.K., Nanda, P.K., Dandapat, P., Bandyopadhyay, S., Gullón, P., Sivaraman, G.K., McClements, D.J., Gullón, B. and Lorenzo, J.M. (2021). Edible Mushrooms as Functional Ingredients for Development of Healthier and More Sustainable Muscle Foods: A Flexitarian Approach. Molecules, 26: 2463.

10. Derbyshire, E.J. and Delange, J. (2021). Fungal Protein—What Is It and What Is the Health Evidence? A Systematic Review Focusing on Mycoprotein. Front. Sustain. Food System, 5: 581682.

11. Dudekula, U.T., Doriya, K. and Devarai, S.K. (2020). A Critical Review on Submerged Production of Mushroom and Their Bioactive Metabolites. Biotechnology, 10:337.

12. Elisashvili, V. (2012). Submerged Cultivation of Medicinal Mushrooms: Bioprocesses and Products. International Journal of Medicinal Mushrooms, 14: 211–239.

13. Jach, M.E., Serefko, A., Ziaja, M. and Kieliszek, M. (2022). Yeast Protein as an Easily Accessible Food Source. Metabolites, 12: 63.

14. Kim, J., Hwang, S. and Lee, S.M. (2022). Metabolic Engineering for the Utilization of Carbohydrate Portions of Lignocellulosic Biomass. Metabolites Engineering, 71: 2–12.

15. Koppram, R., Tomás-Pejó, E., Xiros, C. & Olsson, L. (2014). Lignocellulosic Ethanol Production at High-Gravity: Challenges and Perspectives. Trends Biotechnology, 32, 46–53.

16. Kim, K., Choi, B., Lee, I., Lee, H., Kwon, S., Oh, K. and Kim, A.Y. (2011). Bioproduction of Mushroom Mycelium of Agaricus Bisporus by Commercial Submerged Fermentation for the Production of Meat Analogue. Journal of Science in Food Agriculture, 91: 1561–1568.

17. Papaspyridi, L.M., Aligiannis, N., Topakas, E., Christakopoulos, P., Skaltsounis, A.L. and Fokialakis, N. (2012). Submerged Fermentation of the Edible Mushroom Pleurotus Ostreatus in a Batch Stirred Tank Bioreactor as a Promising Alternative for the Effective Production of Bioactive Metabolites. Molecules, 17: 2714–2724.

18. Pellegrino, R.M., Blasi, F., Angelini, P., Ianni, F., Alabed, H.B.R., Emiliani, C., Venanzoni, R. and Cossignani, L. (2022). LC/MS Q-TOF Metabolomic Investigation of Amino Acids and Dipeptides in Pleurotus Ostreatus Grown on Different Substrates. Journal of Agricultural Food Chemistry, 70: 10371–10382.

19. Ribeiro, B., Andrade, P.B., Silva, B.M., Baptista, P., Seabra, R.M. and Valentão, P. Comparative (2008). Study on Free Amino Acid Composition of Wild Edible Mushroom Species. Journal of Agricultural Food Chemistry, 56:10973–10979.

20. Sydor, M.; Cofta, G.; Doczekalska, B.; Bonenberg, A. (2022). Fungi in Mycelium-Based Composites: Usage and Recommendations. Materials, 15: 6283.

21. Souza Filho, P.F., Nair, R.B., Andersson, D., Lennartsson, P.R., and Taherzadeh, M.J. (2018). Mycoprotein Concentrate from Pea-Processing Industry Byproduct Using Edible Filamentous Fungi. Fungal Biology and Biotechnology, 5: 5.

22. Thiviya, P., Gamage, A., Kapilan, R., Merah, O. and Madhujith, T. (2022). Single Cell Protein Production Using Different Fruit Waste: A Review. Separations, 9: 178.

23. Xiao, Q., Ma, F., Li, Y., Yu, H., Li, C. and Zhang, X. (2017). Differential Proteomic Profiles of Pleurotus ostreatus in Response to Lignocellulosic Components Provide Insights into Divergent Adaptive Mechanisms. Frontier Microbiology, 8: 480.

24. Yan, Z., Zhao, M., Wu, X. and Zhang, J. (2020). Metabolic Response of Pleurotus ostreatus to Continuous Heat Stress. Frontier Microbiology, 10: 3148.

25. Zhao, Z., Xian, M., Liu, M. and Zhao, G. (2020). Biochemical Routes for Uptake and Conversion of Xylose by Microorganisms. Biotechnology and Biofuels, 13: 21.

26. Zikou, E., Chatzifragkou, A., Koutinas, A.A. and Papanikolaou, S. (2023). Evaluating Glucose and Xylose as Cosubstrates for Lipid Accumulation and γ-Linolenic Acid Biosynthesis of Thamnidium Elegans. Journal of Applied Microbiology, 114: 1020–1032..

